# Machine Learning Identifies Ponatinib as a Potent Inhibitor of SARS-CoV2-induced Cytokine Storm

**DOI:** 10.1101/2021.04.07.438871

**Authors:** Marina Chan, Siddharth Vijay, M. Juliana McElrath, Eric C. Holland, Taranjit S Gujral

## Abstract

Although 15-20% of COVID-19 patients experience hyper-inflammation induced by massive cytokine production, cellular triggers of this process and strategies to target them remain poorly understood. Here, we show that the N-terminal domain (NTD) of the spike protein from the SARS-CoV-2 and emerging variants B1.1.7 and B.1.351 substantially induces multiple inflammatory molecules in human monocytes and PBMCs. Further, we identified several protein kinases, including JAK1, EPHA7, IRAK1, MAPK12, and MAP3K8, as essential downstream mediators of NTD-induced cytokine release. Additionally, we found that the FDA-approved, multi-kinase inhibitor Ponatinib is a potent inhibitor of the NTD-mediated cytokine storm. Taken together, we propose that agents targeting multiple kinases required for the SARS-CoV-2-mediated cytokine storm, such as Ponatinib, may represent an attractive therapeutic option for treating moderate to severe COVID-19.

## Main

The severe acute respiratory syndrome coronavirus 2 (SARS-CoV-2) is the viral pathogen responsible for the current Coronavirus Disease 2019 (COVID-19) pandemic. COVID-19 typically presents with symptoms attributed to viral replication that resolve within 1-2 weeks. In approximately 15–20% of cases, this initial phase is followed by more serious events where monocytes produce a cytokine storm with the rapid release of IL-6, IL1b, CXCL10, CCL7 and other inflammatory molecules (*1*). The presence of these cytokines in the patient samples were found to be associated with increased viral load, loss of lung function, lung injury, and a fatal outcome (*2*). Therefore, drugs or drug combinations that reduce the cytokine storm would be useful for treatment of COVID-19. However, although it is known that the induction of the cytokine storm is regulated by and requires IL-1 signaling, the downstream signaling pathways required for this event are not fully identified, thus impeding the development of targeted therapies. Here, we combined approaches from immunology, computational biology, and biochemistry to uncover underlying signaling networks that trigger cytokine storm in monocytes, and we identify and validate FDA-approved or clinical grade compounds that inhibit the production of inflammatory cytokines in the context of SARS-CoV-2 infection. Our goal was to determine if FDA-approved drugs available for clinical use might reduce the time of drug development and deliver a timely solution for the COVID-19 pandemic.

To model monocyte-SARS-CoV-2 interaction *in vitro*, we used the spontaneously immortalized monocyte-like cell line, THP1(*3*), and mammalian (HEK293) cells that produced full-length Spike subunit S1 protein (Val^16^-Gln^690^), which is the SARS-CoV-2 protein essential for host cell entry (**Fig. 1A**). Consistent with previous observations, our preliminary data show that 24-hour stimulation with full-length mammalian cell-derived S1 subunit of SARS-CoV-2 spike protein causes massive upregulation of IL-1b in a dose-dependent manner (**Fig. S1**). Next, we asked if S1 protein could promote upregulated expression of cytokines observed to be elevated in COVID-19 patients. Our data show that S1 protein stimulation (1μg/mL) causes a significant increase in the expression of a panel of cytokines including IL-8 (∼96-fold), IL-6 (5-fold), IL1b (∼120-fold), TNF (∼15-fold), and chemokines including CXCL10 (600-fold), CCL2 (∼70-fold), CCL7 (4-fold) in THP1 monocytes (**Fig. 1B**). Similar changes in cytokines were also observed in healthy donor PBMCs and Raw264.7 mouse monocytes in response to full-length S1 protein (**Fig. S1)**. Together, these data indicate that interaction between Spike subunit S1 protein and monocytes is sufficient to activate monocytes.

**Fig. 1.**
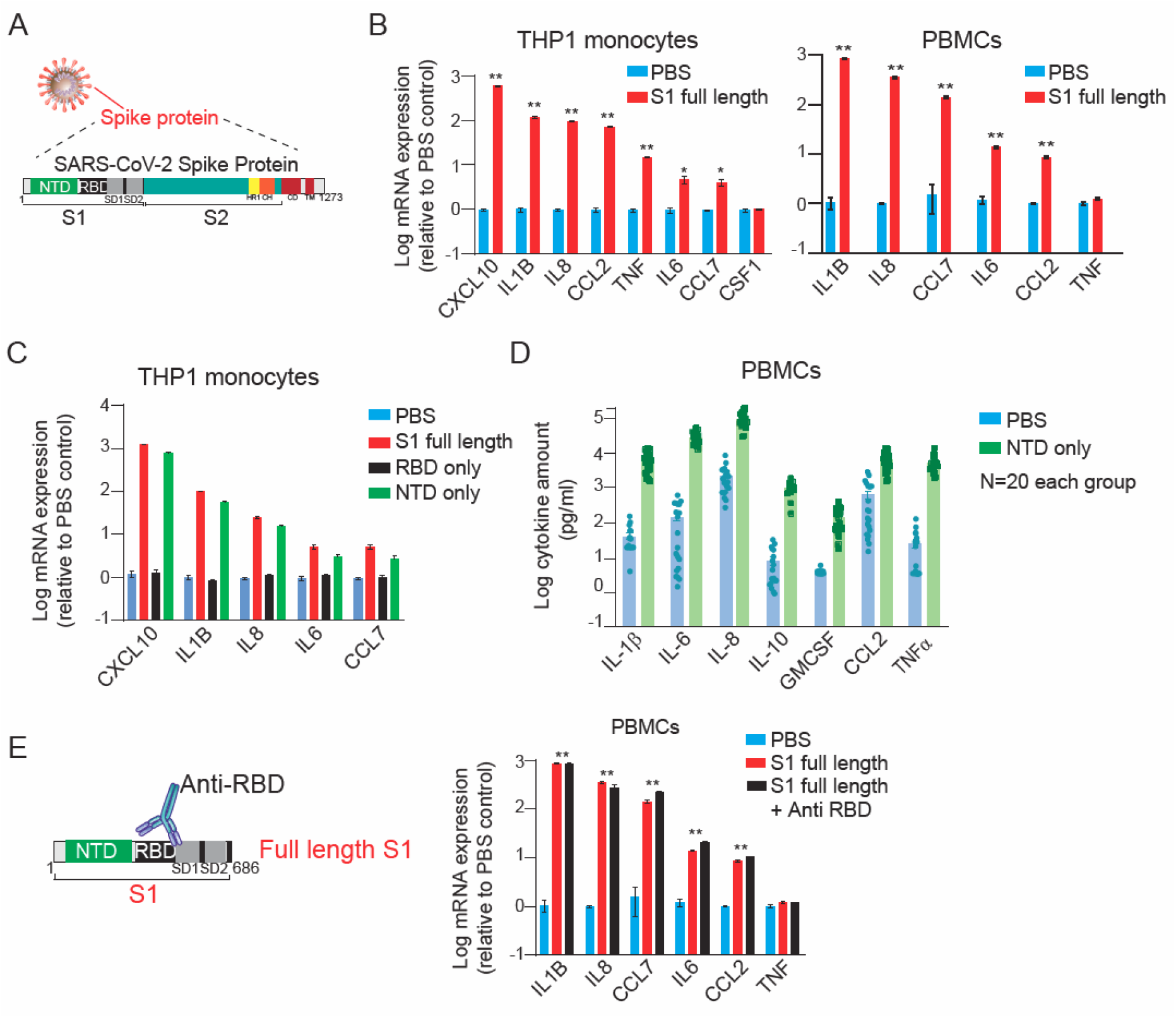
SARS-Cov-2 Spike subunit S1 protein causes a significant increase in the expression and release of a panel of cytokines in THP1 monocytes and human PBMCs. (**A**) A schematic showing major SARS-CoV-2 proteins and domain structure of spike protein. (**B**) Changes in the expression of cytokines in THP1 macrophages (left) and PBMCs (right) upon treatment with full-length S1 subunit at 1 µg/ml for 24 hr. (**C**) Changes in the expression of cytokines in THP1 macrophages upon treatment with different domains of S1 subunit at 1 µg/ml for 24 hr. (**D**) Measurement of cytokine release from healthy donor PBMCs treated with PBS or NTD at 1 µg/ml for 24 hr. (**E**) Effect of an anti-RBD antibody on S1 subunit stimulated changes in the expression of cytokines in PBMCs. Gene expression was measured by qPCR. Cytokine release in the conditioned media was measured by Luminex. Full-length S1 and S1 subunits are purified from HEK293 cells.

Next, we sought to determine the region of the S1 subunit that mediates monocyte activation. It has been established that SARS-CoV-2 and SARS-CoV recognize the angiotensin-converting enzyme 2 (ACE2) receptor in humans (*4*). Specifically, the receptor-binding domain (RBD) of the spike subunit S1 protein of SARS-CoV-2 binds to ACE2 (*4*), thereby promoting the entry of the virus into the cells while the function of the N-terminal domain (NTD) is not well understood. Surprisingly, we found that stimulation with the HEK293 cells-produced NTD (Val^16^- Ser^305^) of the S1 subunit is sufficient to promote cytokine expression while stimulation with the RBD (Arg^319^-Phe^541^) of S1 protein could not activate THP1 monocytes (**Fig. 1C**). Similar changes in the release of cytokines in the conditioned media were also observed in PBMCs from 20 healthy individuals in response to stimulation with NTD of S1 subunit (**Fig. 1D**). Consistently, treatment with CV30(*5*) (high potent antibody targeting RBD of SARS-CoV-2) did not change S1 protein-induced expression of cytokines in PBMCs (**Fig. 1E**). Further, stimulation with spike protein from two different endemic coronaviruses (HCoV-COV-2 and HCoV-OC43) did not promote cytokine release in THP1 cells (**Fig.S1**), suggesting that the S1 subunit of SARS-CoV-2 contains a unique feature of this specific spike protein sequence required for monocytes activation. Together, these data suggest that NTD of S1 subunit interacts with a different receptor than ACE2, on monocytes to activate the ‘cytokine storm.’

To identify signaling pathways activated in monocytes in response to S1 spike protein and explore potential targets for therapeutic development, we employed a recently developed strategy that combines phenotypic screening with machine learning-based functional screening approaches, called KiDNN (*6*) and KiR (*7, 8*). These strategies utilize large-scale drug-target profiling efforts, Regularized regression or Deep Neural Networks (DNN), and broadly selective chemical tool compounds to pinpoint specific nodes (kinases and associated networks) underlying a given phenotype, such as cell growth or release of secreted factors, e.g., cytokines (**Fig. 2A**). We screened a set of 35 computationally-chosen kinase inhibitors (*7*) and quantified their effect on changes in NTD-mediated release of seven cytokines in the conditioned media from pooled PBMCs (**Fig. 2B, Dataset 1**). Using this training dataset, we built both elastic net regularization (**Fig. S2**) and preliminary DNN models (**Fig. S3**) to predict kinases essential for the NTD-mediated release of cytokines. Using a multi-phase grid search, we optimized the hyperparameters for each preliminary DNN model (**Fig. S3**). Model performance was evaluated using leave one out cross-validation (LOOCV) mean squared error (MSE) between predicted and observed drug response. In LOOCV, each time 34 drugs’ activity profiles were used to train the model to predict the remaining drug’s effect on NTD-mediated cytokine release. MSE between predicted and observed cytokine levels was used to assign an error score to each model. Overall, models for each of the cytokines performed with at least 85% accuracy (**Fig. S2**). The optimized models with the least mean squared errors collectively identified 30 most “informative kinases” (out of >300 kinases) that may be involved in the NTD-mediated cytokine release (**Fig. S4**). These include several kinases known to play a critical role in cytokine signaling, such as JAK1 and IRAK1, as well as kinases not previously known to play a role in this process, like EPHA7, MAP3K8, and MAP3K2. Overall, MAP3K8 was shown to be enriched in the cytokine-mediated signaling network of all seven cytokines, while EPHA7 was enriched in networks of 4 out of 7 cytokines (**Fig. S4**). Therefore, based on this analysis, we predicted that both MAP3K8 and EPHA7 are essential for the NTD-mediated release of cytokines.

**Fig. 2.**
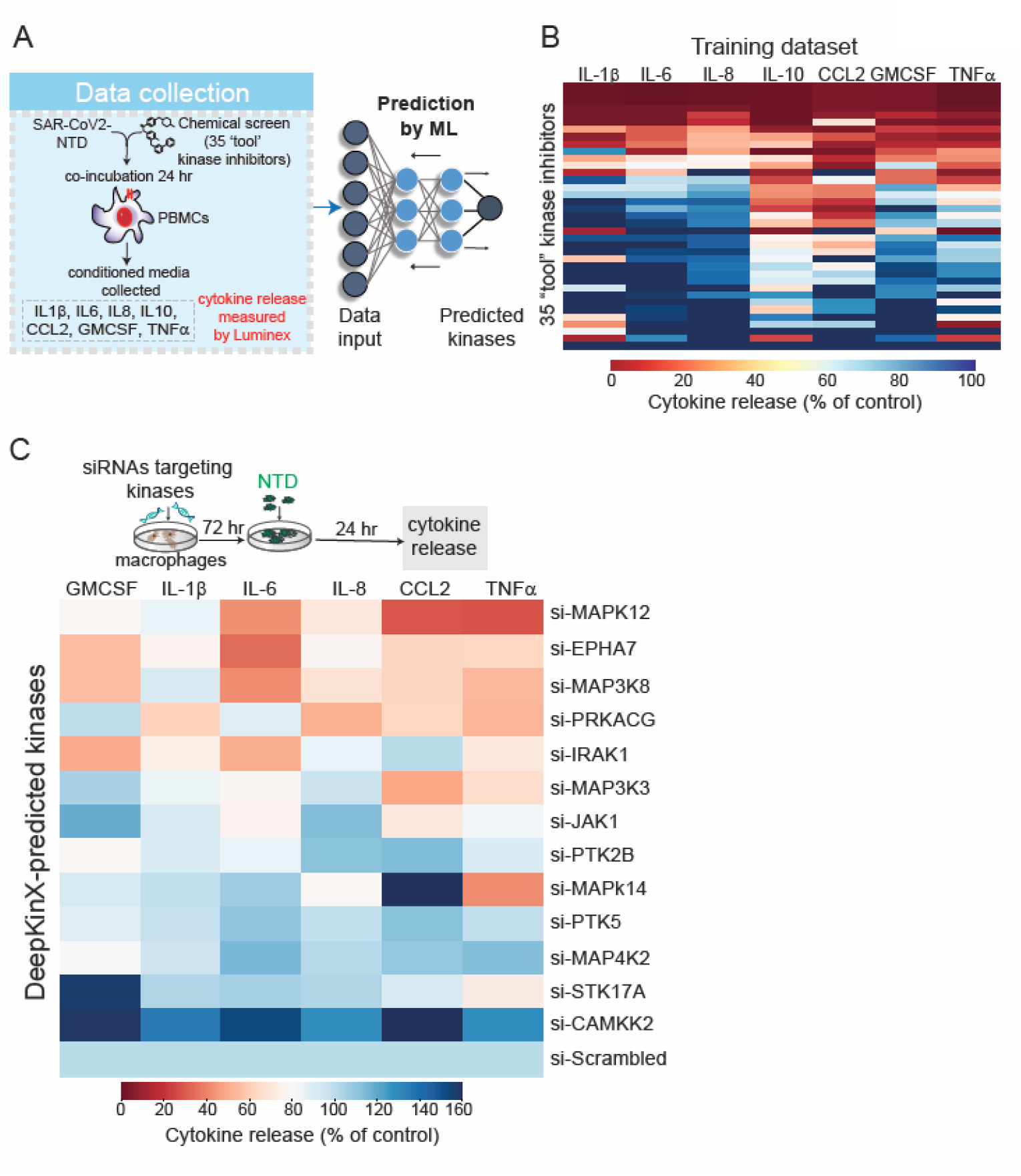
Machine learning-based functional screening identified key kinase drivers of the NTD stimulated cytokine release in PBMCs. (**A**) A schematic showing the process of the machine learning-based functional screening. (**B**) A heatmap showing changes in indicated cytokines in response to 500 nM treatment of PBMCs with 35 “tool” kinase inhibitors as the training set. Cytokines released in the media were measured by Luminex and normalized to DMSO control. (**C**) Validation of predicted kinases as drivers of cytokine release by siRNAs in THP1 cells. Cytokine release was measured by Luminex and normalized to scrambled siRNA.

To validate the role of the kinases that we predicted to be important in cytokine signaling, we examined the effects of depleting these kinases in gene knockdown experiments. Using a pooled set of four siRNA, we knocked down the expression of top 13 kinases implicated by our KiDNN and KiR analyses (**Fig. S4**) in THP1 cells and measured their effect on NTD-mediated cytokine release. Transient transfections of pooled siRNA led to a 35-80% knockdown of each of the kinases measured by quantitative PCR (**Fig. S4**). Our data showed that knockdown of 11 out of 13 kinases led to an >1.5-fold decrease in NTD-mediated cytokine levels compared with scrambled siRNA control (**Fig. 2C**). Of these, knockdown of MAPK12, EPHA7, MAP3K8, PRKACG, IRAK1, MAP3K3, and JAK1 led to a decrease in more than one cytokine or chemokine (**Fig. 2C**). Network analysis based on prior information showed that several of these kinases are known to regulate common transcription factors such as *MYC* and *REL* (**Fig. S4)**. Together, these data confirmed KiR and KiDNN models’ predictions and validated the role of at least seven kinases in the SARS-CoV2 S1 subunit NTD-mediated cytokine storm.

Given that our analysis implicated multiple kinases that affect different signaling pathways in cytokine storm triggering, we hypothesized that a rational drug combination or multi-targeted therapy may represent an effective approach to blocking the SARS-CoV-2 mediated cytokine storm. Therefore, by taking advantage of non-linearity of DNN models and cloud computing, we predicted responses to 91 thousand two-drug combinations, as well as responses to 13 million combinations involving three-drug cocktails. Next, we used the optimized KiDNN models to predict both pairwise and single-agent responses to 427 single inhibitors that reduce NTD-mediated cytokine levels (**Fig. 3A**). In our analysis, we prioritized those compounds that are FDA-approved for human use, known to exhibit a low toxicity profile, and predicted to inhibit the release of multiple cytokines. Combinations containing the most promiscuous drugs, such as staurosporine and K252a, were excluded from this analysis. Our model indicated several FDA-approved compounds, including Ponatinib, Baricitinib, and Combimetinib, could inhibit NTD-mediated cytokine storm to varying levels as a single agent (**Dataset 1**). Of these, Ponatinib, an FDA-approved drug for chronic myelogenous leukemia, was predicted to be the most effective in blocking all seven cytokines as a single agent, as well as in combination (top 360 out of 500 combinations). We experimentally confirmed that treatment with Ponatinib potently inhibits NTD-mediated cytokine release in PBMCs in a dose-dependent manner (EC_50_ 10-28nM) **(Fig. 3B, S5**). No significant changes in the viability of PBMCs were observed, even 1uM Ponatinib treatment (**Dataset 1**). Further, treatment with Ponatinib outperforms Baricitinib, a JAK inhibitor FDA-approved for the treatment of COVID-19 (*9*), in inhibiting all seven cytokines in response to NTD in PBMCs (**Fig. S5**). Kinase activity profiling of Ponatinib showed that this drug inhibits 8 out of 10 experimentally-validated kinases necessary for the NTD-mediated cytokine and chemokine release (**Fig. 3C, D**). In contrast, Baricitinib was shown to inhibit only 4 of these essential kinases. Thus, our data indicate that Ponatinib, a multi-specific kinase inhibitor, blocks the activity of several kinases that are essential for cytokine signaling thereby dampening the NTD-mediated cytokine storm.

**Fig. 3.**
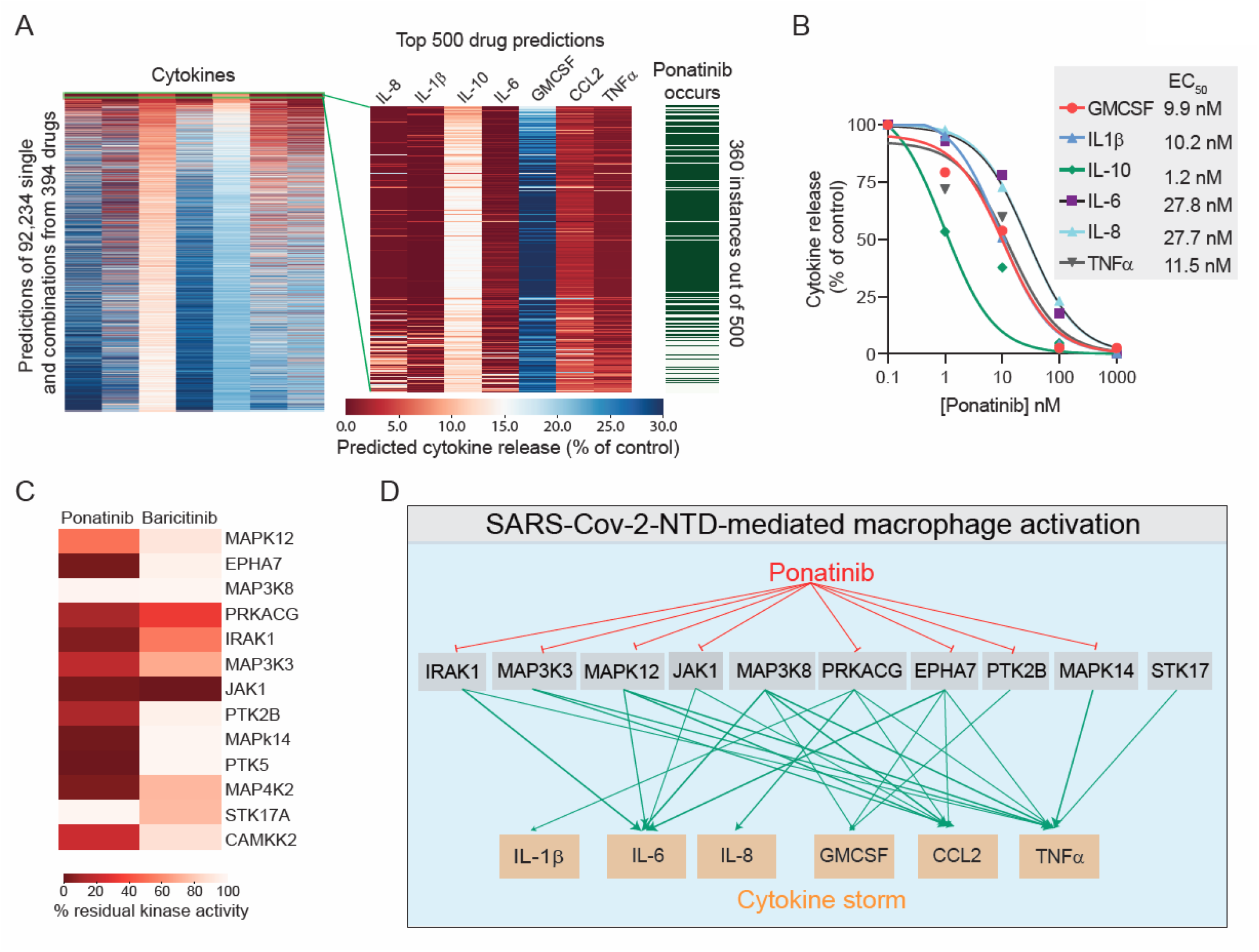
Machine learning-based functional screening identified Ponatinib as a potent inhibitor of NTD-mediated cytokine storm. (**A**) A heatmap showing the effect of single and combinatorial drug predictions by machine learning on indicated cytokine release (left), a zoomed view of top 500 drug predictions (middle), and frequency of Ponatinib occurrence in the top 500 predictions (right). A total of 92,234 single and combinatorial drug predictions were generated from 394 drugs. Predicted cytokine release is expressed as % of DMSO control. (**B**) Dose-response curves of ponatinib treatment on cytokine release in PBMCs. (**C**) Comparison of kinase inhibition profile of ponatinib and baricitinib. (**D**) A schematic showing ponatinib inhibits multiple kinases involved in the SARS-CoV2-NTD-mediated cytokine signaling.

Recently, several new variants of SARS-CoV-2 have emerged, including the B.1.1.7 first identified in the United Kingdom and B.1.351 identified in South Africa that harbor mutation sites in the NTD. We asked if these variants could also promote cytokine release. We show that both B1.1.7 and B.1.351 NTDs could promote the release of cytokines in PBMCs at the levels comparable with the NTD from Wuhan SARS-CoV-2 (**Fig. 4A**). Notably, Ponatinib treatment could inhibit the NTD-mediated release of cytokines from all variants. Further, Ponatinib treatment outperforms both dexamethasone and Baricitinib to block the cytokine release at the same dose (**Fig. 4A**). Finally, we evaluated the response of Ponatinib in PBMCs from 19 COVID19 patients. Our data show that NTD stimulation (1μg/mL) causes a significant increase in the release of all seven cytokines measured in the conditioned media from COVID19 PBMCs (**Fig. 4B**). Interestingly, all cytokines and chemokines showed a large variable response to NTD, suggesting an inherent patient-to-patient variability. However, treatment with 1uM Ponatinib completely shuts down NTD-mediated cytokine storm in all COVID-19 PBMCs. In contrast, 1uM dexamethasone, a corticosteroid prescribed to COVID-19 patients, showed a weaker response in the presence of NTD. Overall, these data provide a strong rationale for repositioning Ponatinib to dampen the SARS-CoV-2-induced cytokine storm and be immediately tested in the clinic.

**Fig. 4.**
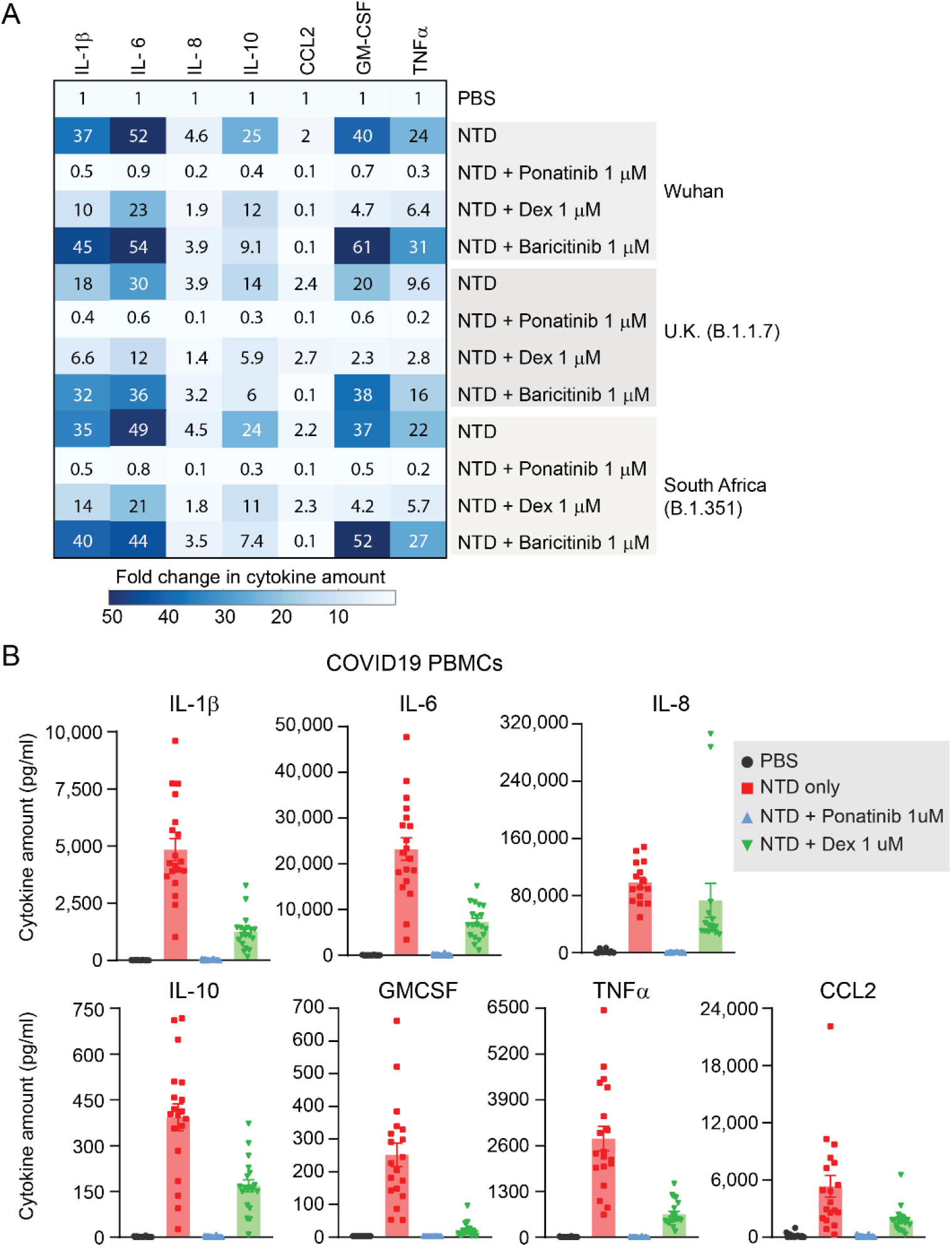
Ponatinib inhibits NTD-mediated cytokine storm in PBMCs from COVID19 patients. **(A)** A heatmap showing changes in indicated cytokines in response to NTD from indicated SARS-Cov-2 variants and inhibitors. Data are shown as fold change between NTD treatment and PBS control. Dex; dexamethasone. **(B)** Plots showing changes in indicated cytokines in response to NTD at 1 µg/ml and ponatinib or dexamethasone in COVID19 PBMCs for 24 hours. Cytokines released in the media were measured by Luminex and normalized to DMSO control. Bars represent the mean of 19 individual donors. Error bars represent SEM.

In summary, we discovered a previously unknown function of the NTD of the SARS-CoV-2 spike protein in promoting cytokine release in immune cells. We identified several new host-specific kinases that are required for the SARS-CoV-2 spike protein-mediated release of cytokines and chemokines in monocytes. Our findings strongly suggest that simultaneous targeting of multiple host kinases involved in SARS-CoV-2 mediating cytokine storm yield more effective treatment options than the use of more selective agents. More specifically, our results suggest that an FDA-approved drug, Ponatinib, could represent a strong candidate for drug repurposing efforts aimed at providing an alternative, and timely treatment for COVID-19 patients exhibiting major, life-threatening symptoms.

## Acknowledgments

This work was supported by grants from the Fred Hutch COVID19 Pilot Fund and Scientific Computing Infrastructure at Fred Hutch funded by ORIP grant S10OD028685. We thank Dr. Milka Kostic for her helpful comments and suggestions for the manuscript.

## Supplementary Information

Supplementary Figures S1-S5, Dataset 1

Online Materials and Methods

**Figure S1.**
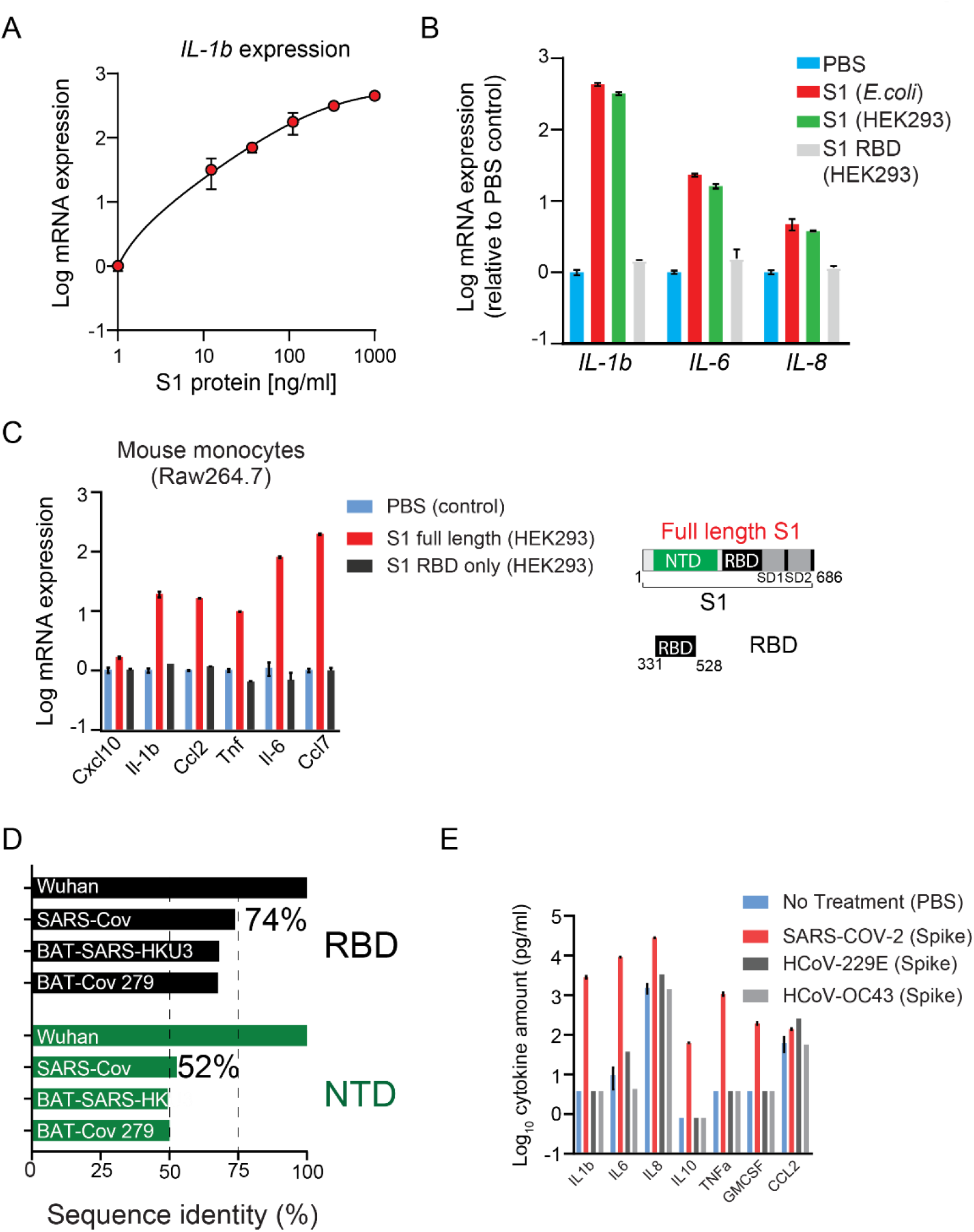
SARS-Cov-2 Spike subunit S1 protein causes a significant increase in the expression of a panel of cytokines in monocytes. (A) Plot showing dose-dependent increase in *IL1b* expression in THP1 cells in response to S1 protein. (B) S1 RBD domain does not activate THP1 monocytes. THP1 cells were stimulated with indicated protein (1μg/mL) or PBS for 24 hours. Gene expression was measured by qPCR. (C) SARS-Cov-2 spike protein activates mouse macrophages *in vitro*. Plots showing changes in the expression of cytokines in mouse Raw264.7 macrophages upon treatment with different domains of S1 subunit at 1 µg/ml for 24 hr. (D) The NTD is more divergent than the RBD of SARS-Cov-2. Plot shows sequence homology of NTD and RBD region of the SARS-Cov-2 spike protein with other coronaviruses. (E) Spike protein from endemic viruses does not activate THP1 monocytes. THP1 cells were stimulated with indicated protein (1μg/mL) or PBS for 24 hours. Gene expression was measured by qPCR.

**Figure S2.**
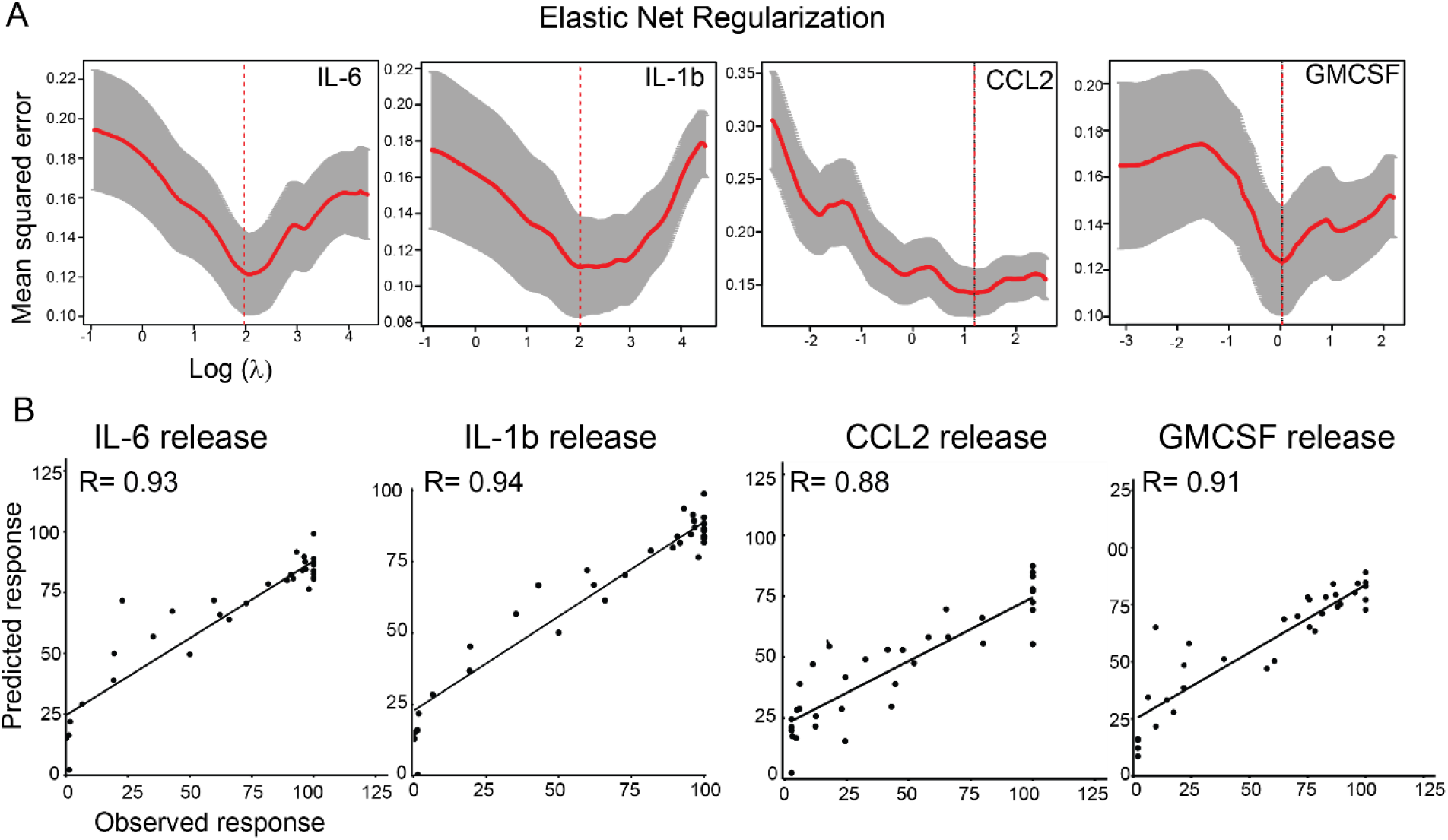
Applying kinase inhibitor regularization (KiR) to SARS-Cov-2 Spike subunit S1 mediated changes in cytokine release. (A) Representative plots showing leave-one-out-cross validation error using elastic net regularization fit for indicated cytokines. The error bars (grey) represent cross-validation error plus 1SD. The kinases identified at absolute minima (red dashed line) were termed the most informative kinases. (B) Plot showing correlation between model-predicted and observed response in pooled PBMCs for indicated cytokines.

**Figure S3.**
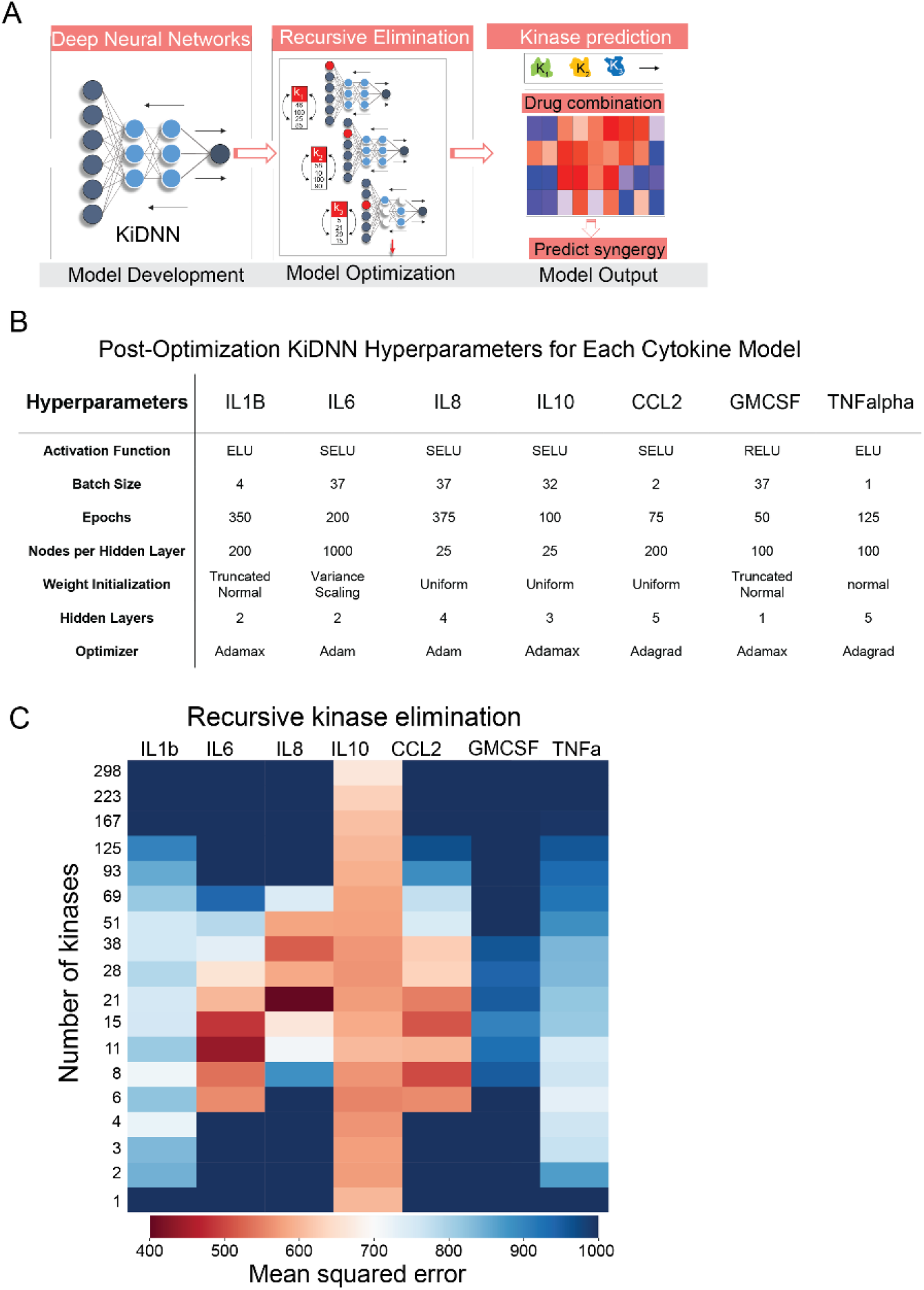
KiDNN Optimization. (A) A schematic illustrating the KiDNN modeling was used to identify underlying essential kinases and kinase inhibitor combinations that could inhibit NTD-mediated cytokine release. (B) A table showing a set of hyperparameters used for developing KiDNN models for each cytokine. ReLU; rectified linear unit, ELU; exponential linear unit, SeLU; Scaled Exponential Linear Unit, Adagrad; adaptive gradient. (B) A heatmap showing mean squared error following the recursive kinase elimination step for each KiDNN model.

**Figure S4.**
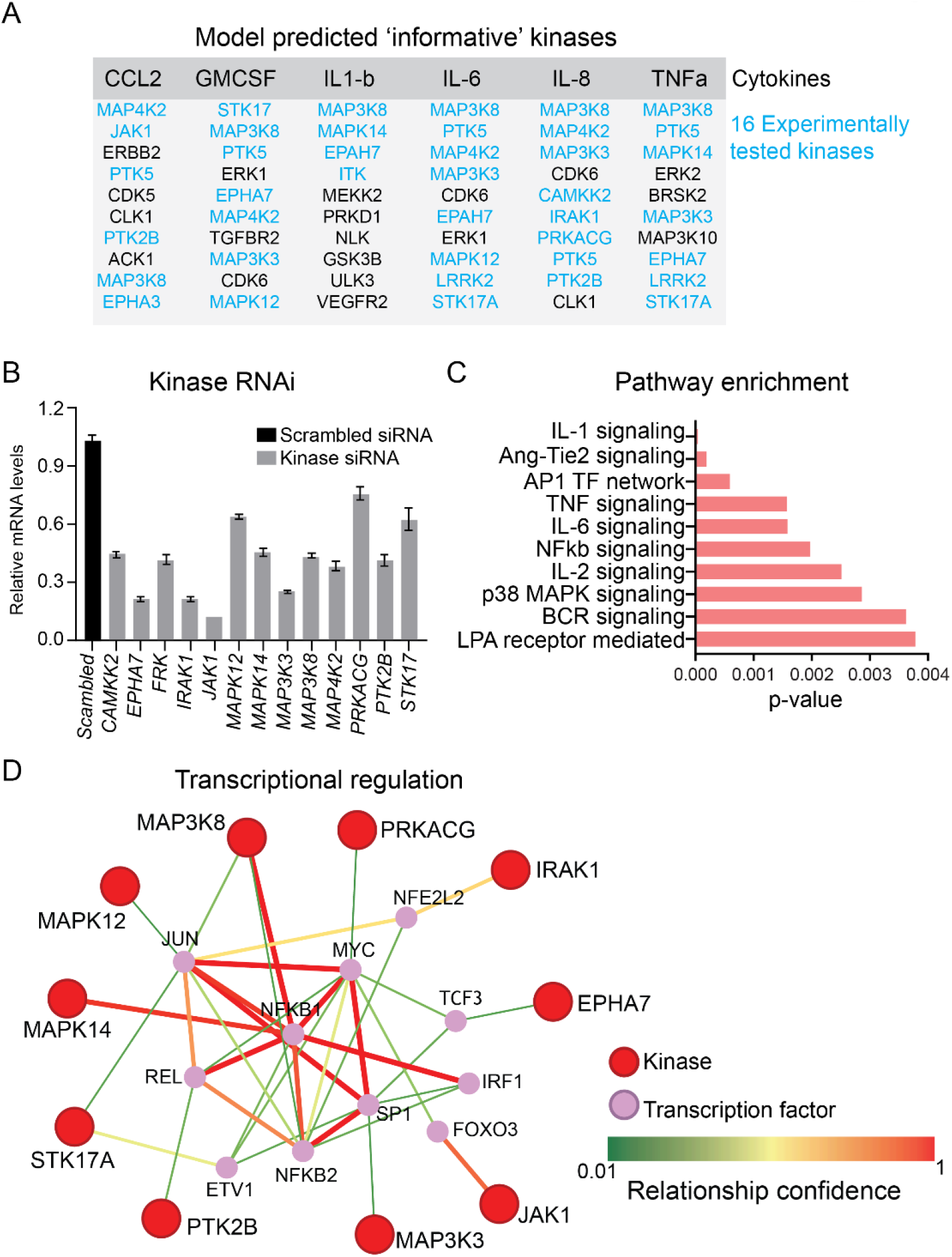
Model-predicted informative kinases. (A) A table showing list of kinases predicted to be necessary for the release of indicated cytokines by both KiR and KiDNN modeling. Sixteen kinases from this list were chosen for experimental validation (shown in blue). (B). Plot showing knockdown efficiency of siRNA targeting indicated kinases compared with scrambled control. Data are presented as means of three replicates, error bars represent SEM. (C) Pathway enrichment plot generated using PathwayNet^1^ using input from model-predicted kinases. (D) Schematic showing known interactions between model-predicted kinases and transcription factors generated using PathwayNet^1^.

**Figure S5.**
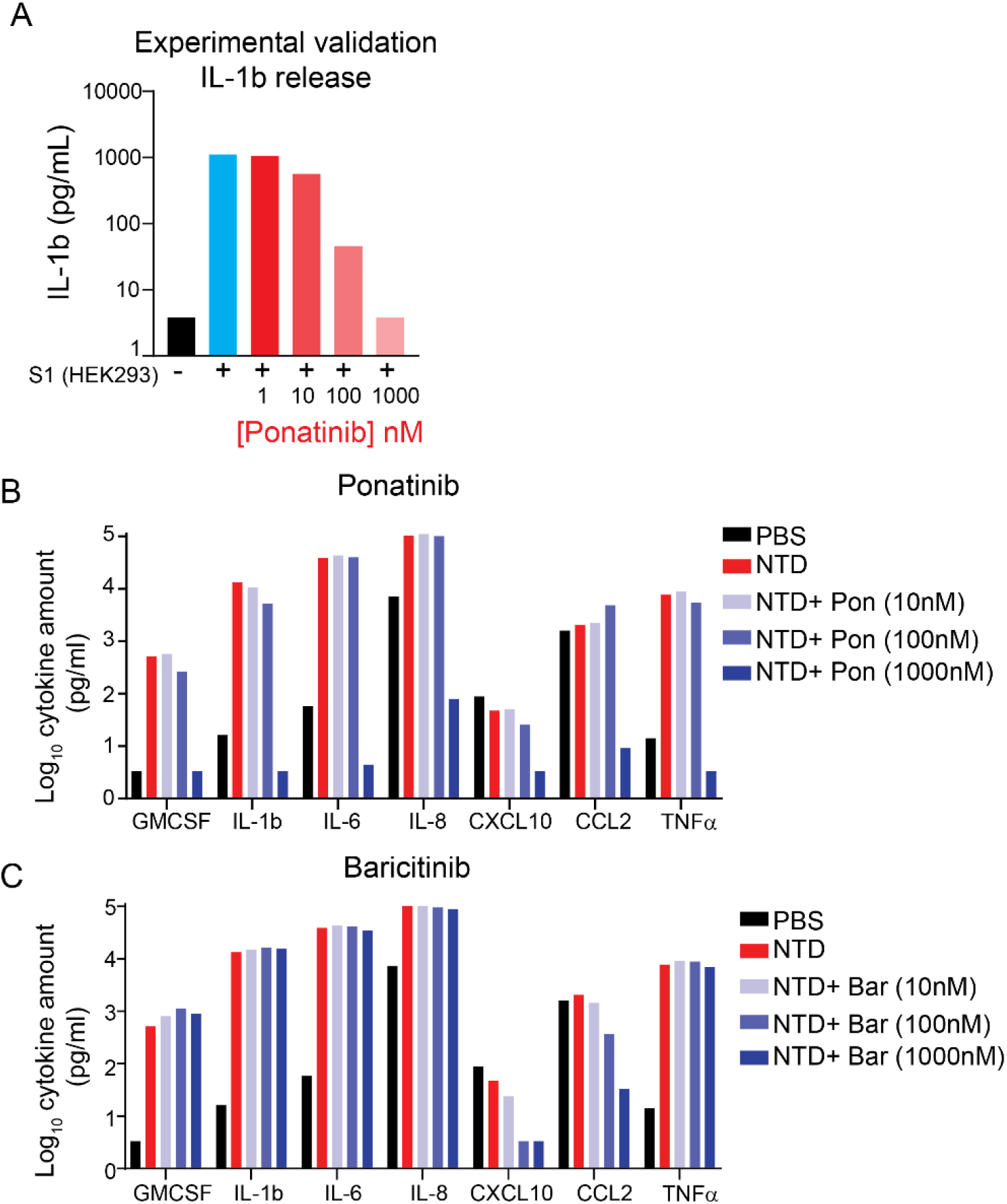
Ponatinib inhibits NTD-mediated cytokine storm in PBMCs. (A) Plot showing Ponatinib (the top model-predicted, FDA-approved) treatment significantly decreases S1-mediated IL-1b release in a dose-dependent manner. (B, C) Plots showing changed in the nTD-mediated release of indicated cytokines in response to Ponatinib (B) and Baricitinib (C) at indicated doses. Cytokines released in the conditioned media from PBMCs were measured using Luminex.

## Methods

Methods, including statements of data availability and any associated accession codes and references, are available in the online version of the paper.

### Cell lines, chemicals and purified proteins

THP1 and Raw264.7 cells were obtained from American Type Culture Collection. Peripheral blood mononuclear cells (PBMCs) from healthy donors spanning various age groups were obtained from Bloodworks NW, Seattle, Washington. PBMCs and THP-1 cells were cultured in RPMI1640 media supplemented with 10% FBS, 1% P/S, 1 mM sodium pyruvate. Raw264.7 cells were maintained in Dulbecco’s minimum essential medium supplemented with 10% FBS (Sigma) and 1% Penn Strep. All cell lines were grown at 37°C under 5% CO2, 95% ambient atmosphere. Ponatinib (Cat# S1490), dexamethasone (Cat# S1322) and baricitinib (Cat# S2851) were obtained from Selleck Chemicals. Recombinant purified full length S1 protein (Cat# 230-30161), and the receptor biding domain (RBD; Cat # 230-30162) from SARS-CoV-2 were obtained from Ray Biotech Life. Recombinant purified N-terminal domain (NTD) of the SARS-CoV-2 was obtained from Leinco Technologies Inc (Cat #S853) or Acro Biosystems (Cat # S1D-C52H6). Spike protein from HCoV-229E (Cat# 40605-V08B) and HCoV-OC43 (Cat# 40607-V08H1) were obtained from Sino Biological Inc. Recombinant purified NTD of the SARS-CoV-2 emerging variants were obtained from Acro Biosystems (Cat # S1D-C52H9 and # S1D-C52H7).

### Human peripheral blood mononuclear cells from COVID19 patients

Consenting SARS-CoV-2-infected (n=19) donors, age 18 years and older, provided anticoagulated blood samples by venipuncture at the Seattle Vaccine Trials Unit. SARS-CoV-2 donors were diagnosed by PCR testing of nasopharyngeal swabs, had mild-moderate disease and were sampled post-symptom onset. PBMC were isolated and cryopreserved within four hours of collection. Cell viabilities were assessed post-thawing and after 24 hours of treatment. Fred Hutchinson Cancer Research Center Institutional Review Board approved all aspects of this study (IRB 10440, 00001080 and 00022371).

### Cytokine Measurement

Cytokines were measured by Luminex multiplex assay. Samples and cytokine standards were incubated with Luminex microbeads (one unique bead population per cytokine) coated with cytokine-specific antibodies. Beads are washed then incubated 1 hour with biotinylated anti-cytokine antibodies, washed again then incubated 30 minutes with a phycoerythrin-streptavidin conjugate. After a final wash the assay is read on a Luminex 200 instrument, classifying each bead as to its cytokine-specificity and phycoerythrin fluorescence intensity. Phycoerythrin fluorescence of each bead will be proportional to the cytokine concentration in the samples or standards. A 5-parameter logistic standard curve is generated for each cytokine, with sample concentrations calculated from these curves.

### Kinase inhibitor screening

Kinase inhibitor screening was performed as described previously^2^. Briefly, 35 kinase inhibitors were tested for the effect on NTD-mediated cytokine release in PBMC. All inhibitors were tested at 500nM. Pooled PBMC from several donors were plated in 12-well plate (1×10^6^ cell per well in 1mL). Kinase inhibitors or DMSO control was subsequently added to each well. Conditioned medium collected 24 hours post treatment was snap frozen for cytokine analysis.

### Cell viability

The effects of inhibitors as single agent on viability of PBMCs was measured using CellTiter-Glo assay (Promega, WI, USA) as described previously^3, 4^. Briefly, cells (5×10^3^ in 100 ul culture medium) were seeded on a 96-well plate (Corning, NY, USA). Cells were then treated with various inhibitors at 500nM as single agent. After 24 hours, cells were incubated with CTG2.0 reagent for 5 min and total viability was measured by obtaining luminescent signal intensity. The quantified data was normalized to untreated control and plotted in Prism (Graphpad software, San Diego, CA, USA).

### Small interfering RNA transfection

All small interfering RNA (siRNA) targeting human kinases were obtained from Dharmacon (Thermo). siRNA’s transfections in 12-well plate for expression and cytokine profiling was carried out using Lipofectamine RNAiMax (Invitrogen) according to manufacturer instructions.

### RNA extraction and quantitative PCR

Total cellular RNA was isolated using an RNeasy Mini Kit (QIAGEN). mRNA expression changes in genes encoding for various cytokines wer determined using quantitative real-time PCR (qPCR). Briefly, 0.5-1 μg of total RNA was reverse transcribed into first-strand cDNA using an RT2 First Strand Kit (QIAGEN). The resultant cDNA was subjected to qPCR using human cytokine-specific primer (Realtimeprimers.com) and *GAPDH* (housekeeping control). The qPCR reaction was performed with an initial denaturation step of 10 min at 95 °C, followed by 15 s at 95 °C and 60 s at 58 °C for 40 cycles using Biorad CFX384 thermocycler (Biorad). The mRNA levels of genes encoding cytokine expression were normalized relative to the mean levels of the housekeeping gene and compared using the 2−ΔΔCt method as described previously^2^. Primersets to measure cytokine expression were obtained from Realtimeprimers.com.

### Kinase inhibitor Regularization (KiR) modeling

KiR models for NTD-mediated release of each cytokine in PBMCs were generated as previously described (Gujral et al., 2014a). Briefly, a panel of 427 kinase inhibitors previously had their pairwise effects on 298 human kinases profiled (Anastassiadis et al., 2011; Rata et al.). The result is a quantitative drug-target matrix, where each entry is a percentage between 0 and 100 that represents that kinases residual activity (as a percent of control, uninhibited activity) in the presence of that inhibitor. A set of 35 inhibitors were tested on pooled PBMs as described above, with the end result being a single response for each drug that represents the change in cytokine release (as % DMSO control) at the profiled dose of the inhibitor (usually 500 nM). The kinase inhibition profiles of each inhibitor and the quantitative responses to those inhibitors were used as the explanatory and response variables, respectively, for elastic net regularized multiple linear regression models (Zou and Hastie, 2005). Custom R scripts (available at https://github.com/FredHutch/KiRNet-Public) employing the glmnet package were used to generate the final models (Friedman et al., 2010). Leave-one-out cross validation (LOOCV) was used to select the optimal value for the penalty scaling factor λ. Models were computed for 11 evenly-spaced values of α (the relative weighting between LASSO and Ridge regularization) ranging from 0 to 1.0 inclusive. Kinases with positive coefficients in at least one of these models (with the exception of ***α*** = **0**, which always has non-zero coefficients for every kinase) were considered hits (**Figure S4A**). Model accuracy was assessed via the LOOCV error as well as the root-mean-squared error of the predictions for the tested inhibitors (**Figure S2**).

### Deep Neural Network (DNN) Development

The development of the DeepKinX DNN models were achieved through the Keras and TensorFlow Deep Learning framework as described previously^5-7^. Briefly, a multi-phase Grid Search method was used to optimize the DNN hyperparameters (epochs, batch size, optimizer, weight initializer, activation function, hidden layer quantity and nodes per hidden layer)^8^. Grid Search is a commonly employed method of hyperparameter optimization that evaluates combinations of numerous hyperparameter values to identify the model characteristics resulting in the lowest error between observed and predicted migration. The error function that was used to compare numerous models was LOOCV (Leave-One-Out-Cross-Validation) MSE^9^. In LOOCV, each time *n* − 1 drugs’ activity profiles are used to train the model to predict the remaining drug’s effect on cell migration. The process is repeated *n* times, excluding and predicting each and every drug. Mean Squared Error (MSE) between predicted and observed migration is used to assign an error score to each model built with various combinations of hyperparameter values. In each phase of Grid Search, various combinations of hyperparameters are tested and the combination with the lowest LOOCV MSE is used in the subsequent phase of optimization until the final phase is reached. After optimization, the top performing hyperparameters are used to build the DeepKinX network.

### Recursive Kinase Elimination (Mathematical Method)

Assuming a trained DeepKinX DNN model defined by *f* and a dataset defined by [*y, X*_1_, *X*_2_, *X*_3_, …], the baseline error (*e*_*baseline*_) can be estimated, assuming a pre-defined cost function. For both DeepKinX-Fzd2 and DeepKinX- WT models, the cost function was defined as follows:

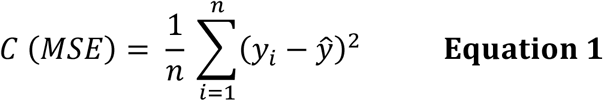

The baseline error was subsequently calculated as defined below:

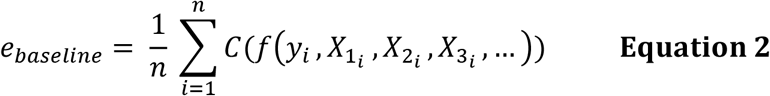

To calculate the post-permutation error (*e*_*permutation*_), each feature is shuffled one-by-one for a total of 10,000 random shuffles. The matrix of features with a single feature permuted once can defined by *X*^*permutation*^. Accordingly, the post-permutation error for an individual feature is computed as follows:

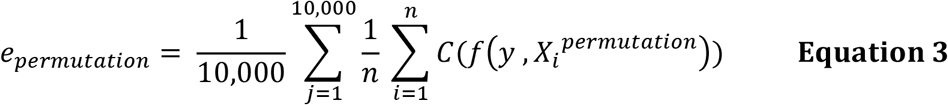

Lastly, the relative kinase importance (*RKI*) or error difference for an individual feature is computed:

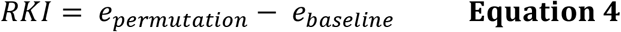

Each kinase is then assigned an RKI score and ranked based from highest to lowest. Subsequently, the bottom 25% of kinases are removed in future iterations of recursive kinase elimination. Using just the top 75% of kinases, a new DeepKinX model is built and LOOCV MSE is used to track the model’s overall relative performance across several rounds. This three-step process— (1) ranking kinases by importance score, (2) removing the bottom 25% of kinases, and (3) assessing LOOCV MSE of the DeepKinX model built using only the remaining kinases— is repeated until the LOOCV MSE of the model reaches an inflection point and starts to increase as the number of inputs decrease.

### Prediction of naïve drugs and drug combinations

To predict the effect of all 427 drugs of the original matrix, the final DeepKinX-Huh7 model was trained on the 44-drug training matrix. Keeping the weights and biases of the DeepKinX-Huh7 network constant, new kinase activity data for naïve drugs were inputted into the model for prediction. Further, the effect of combinations of drugs were also predicted by creating pseudo-activity matrices. Assuming two activity matrices for two different drugs defined by [*J*_1_, *J*_2_, *J*_3_, … *J*_*n*_] and [*K*_1_, *K*_2_, *K*_3_, … *K*_*n*_], where *n* corresponds to a specific kinase’s inhibition, a linear combination of the two activities corresponding to each drug for each kinase was applied to create a pseudo-activity matrix (*P*) combining both drug’s effect on a kinase:

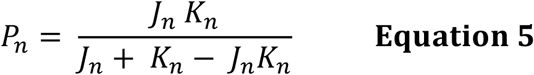

Subsequently, the pseudo-matrix for all 428 by 428 (including control) combinations of drugs was computed and inputted into DeepKinX for prediction. Because combinations of drugs that are effective in combination but not as effective individually are of particular interest, the top 15 drugs predicted individually are removed from the rank-ordered list of predicted viability of all drug combinations. The process of pseudo-matrix creation and successive prediction was similarly extended to 3 drug combos, in which a linear combination of all 3 residual kinase activities for each of 3 drugs was used.

## Data and Code availability

The custom python scripts that implements KiDNN framework are available on GitHub: https://github.com/gujrallab/Covid-19. This repository also includes all associated files needed to execute the script and produce a sample model using the training dataset. R scripts for KiR modeling is available at https://github.com/FredHutch/KiRNet-Public

**Dataset 1. Drug screen data, and machine learning-based predicted kinases and kinase inhibitors**

